# Putting perception into action: Inverse optimal control for continuous psychophysics

**DOI:** 10.1101/2021.12.23.473976

**Authors:** Dominik Straub, Constantin A. Rothkopf

**Affiliations:** Centre for Cognitive Science, Technical University Darmstadt, 64283 Darmstadt, Germany; Institute of Psychology, Technical University Darmstadt, 64283 Darmstadt, Germany; Frankfurt Institute for Advanced Studies, Goethe University Frankfurt, 60438 Frankfurt, Germany

**Keywords:** continuous psychophysics, optimal control, perception and action, inverse reinforcement learning, rational analysis

## Abstract

Psychophysical methods are a cornerstone of psychology, cognitive science, and neuroscience where they have been used to quantify behavior and its neural correlates for a vast range of mental phenomena. Their power derives from the combination of controlled experiments and rigorous analysis through signal detection theory. Unfortunately, they require many tedious trials and preferably highly trained participants. A recently developed approach, continuous psychophysics, promises to transform the field by abandoning the rigid trial structure involving binary responses and replacing it with continuous behavioral adjustments to dynamic stimuli. However, what has precluded wide adoption of this approach is that current analysis methods recover perceptual thresholds, which are one order of magnitude larger compared to equivalent traditional psychophysical experiments. Here we introduce a computational analysis framework for continuous psychophysics based on Bayesian inverse optimal control. We show via simulations and on previously published data that this not only recovers the perceptual thresholds but additionally estimates subjects’ action variability, internal behavioral costs, and subjective beliefs about the experimental stimulus dynamics. Taken together, we provide further evidence for the importance of including acting uncertainties, subjective beliefs, and, crucially, the intrinsic costs of behavior, even in experiments seemingly only investigating perception.

## 1 Introduction

Psychophysical methods such as forced-choice tasks are widely used in psychology, cognitive science, neuroscience, and behavioral economics because they provide precise and reliable quantitative measurements of the relationship between external stimuli and internal sensations (Gescheider, 1997; Wichmann & Jäkel, 2018). The tasks employed by traditional psychophysics are typically characterized by a succession of hundreds of trials in which stimuli are presented briefly and the subject responds with a binary decision, e.g. whether target stimuli at different contrast levels were perceived to be present or absent. The analysis of such experimental data using signal detection theory (SDT; Green & Swets, 1966) employs a model, which assumes that a sensory signal generates an internal representation corrupted by Gaussian variability. A putative comparison of this stochastic signal with an internal criterion leads to a binary decision. These assumptions render human response rates amenable to analysis based on Bayesian decision theory, by probabilistically inverting the model of the decision generating-process. This provides psychologically interpretable measures of perceptual uncertainty and of the decision criterion, e.g. for contrast detection. Thus, the power of classical psychophysics derives from the combination of controlled experimental paradigms with computational analysis encapsulated in SDT.

Psychophysical experiments and their analysis with SDT have revealed invaluable knowledge about perceptual and cognitive abilities, their neuronal underpinnings, and found widespread application to tasks as diverse as eye-witness identification and medical diagnosis (for recent reviews see e.g. Lynn & Barrett, 2014; Wixted, 2020). One central drawback, however, is that collecting data in such tasks is often tedious, as famously noted already by William James (James, 1890). This leads to participants’ engagement levels being low, particularly in untrained subjects, resulting in measurements contaminated by additional variability (Manning et al., 2018). A common solution is to rely on few but highly trained observers to achieve consistent measurements (Green & Swets, 1966; Jäkel & Wichmann, 2006).

Recent work has suggested overcoming this shortcoming by abandoning the rigid structure imposed by independent trials and instead eliciting continuous behavioral adjustments to dynamic stimuli (Bonnen et al., 2015; Bonnen et al., 2017; Huk et al., 2018; Knöll et al., 2018). In their first study using continuous psychophysics, Bonnen et al. (2015) used a tracking task, in which subjects moved a computer mouse to track targets of different contrasts moving according to a random walk governed by linear dynamics with additive Gaussian noise. This experimental protocol not only requires orders of magnitude less time compared to traditional psychophysical methods, but participants report it to be more natural, making it more suitable for untrained subjects. Similar approaches have recently been used to measure contrast sensitivity (Mooney et al., 2018), eye movements towards optic flow (Chow et al., 2021) or retinal sensitivity (Grillini et al., 2021). To obtain measures of perceptual uncertainty from the tracking data, Bonnen et al. (2015) proposed analyzing these data with the generalization of SDT to sequential observations. The Bayes-optimal estimator, in this case, is the Kalman Filter (KF; Kalman, 1960). The KF, however, models the perceptual side of the tracking task only, i.e. how an ideal observer (Geisler, 1989) sequentially computes an estimate of the target’s position. Unfortunately, although the perceptual thresholds estimated with the KF from the tracking task were highly correlated with the perceptual uncertainties obtained with an equivalent classical forced-choice task employing stimuli with the same contrast, Bonnen et al. (2015) reported that these were larger by approximately one order of magnitude.

This is not surprising, since tracking is not merely a perceptual task. In addition to the problem of estimating the current position of the target, a tracking task encompasses the motor control problem of moving the finger, computer mouse, or gaze towards the target. This introduces additional sources of variability and bias. First, repeated movements towards a target exhibit variability (Faisal et al., 2008), which arises because of neural variability during execution of movements (Jones et al., 2002) or their preparation (Churchland et al., 2006). Second, a subject might trade off the instructed behavioral goal of the tracking experiment with subjective costs, such as biomechanical energy expenditure (Di Prampero, 1981) or mental effort (Shenhav et al., 2017). Third, subjects might have mistaken assumptions about the statistics of the task (Beck et al., 2012; Petzschner & Glasauer, 2011), which can lead to different behavior from a model which perfectly knows the task structure, i.e. ideal observers. A model, which only considers the perceptual side of the task, will therefore tend to overestimate perceptual uncertainty, because these additional factors get lumped into perceptual model parameters, as we will show in simulations.

Here, we account for these factors by applying ideas from optimal control under uncertainty (see e.g. Todorov & Jordan, 2002; Wolpert & Ghahramani, 2000, for reviews in the context of sensorimotor neuroscience) to the computational analysis of continuous psychophysics. Partially observable Markov decision processes (POMDPs; Åström, 1965; Kaelbling et al., 1998) offer a general framework for modeling sequential perception and actions under sensory uncertainty, action variability, and explicitly include behavioral costs (Hoppe & Rothkopf, 2019). They can be seen as a generalization of Bayesian decision theory, on which classical analysis of psychophysics including SDT is based, to sequential tasks. Specifically, we employ the well-studied linear quadratic Gaussian setting (LQG; B. D. Anderson & Moore, 2007), which accommodates continuous states and actions under linear dynamics and Gaussian variability. Additionally, the model involves explicit costs, which can accommodate task goals as well as physiological and cognitive costs of performing actions. Finally, we extend the model by allowing subjects to have subjective beliefs about the stimulus dynamics. In the spirit of rational analysis (J. R. Anderson, 1991; Gershman et al., 2015; Simon, 1955) we subsequently invert this model of human behavior, by developing a method for performing Bayesian inference of parameters describing the subject. Importantly, inversion of the model allows all parameters to be inferred from behavioral data and does not presuppose their value, so that, e.g., if subjects’ actions were not influenced by subjective internal behavioral costs, the internal cost parameter would be estimated to be zero. We show through simulations with synthetic data that models, which do not account for behavioral cost and action variability, overestimate perceptual uncertainty when applied to data that includes these factors. We apply our method to data from three previously published experiments, infer perceptual uncertainty, and obtain overwhelming evidence through Bayesian model comparison with the widely applicable information criterion (WAIC) that this model explains the data better than models only considering perceptual contributions to behavioral data. Additionally, the method provides inferences of subjects’ action variability, subjective behavioral costs, and subjective beliefs about stimulus dynamics. Taken together, the methodology presented here improves current analyses and should be the preferred analysis technique for continuous psychophysics paradigms.

## 2 Results

### 2.1 Computational models of psychophysical tasks

In a classical psychophysics task (e.g. position discrimination; Figure 1 A, B), the stimuli *x_t_* presented to the observer are usually independent and identically distributed between trials. This allows for straightforward application of Bayesian decision theory: in any trial, the observer receives a stochastic measurement *y_t_* ~ *p*(*y_t_*|*x_t_*, ***θ***), forms a belief *p*(*x_t_*|*y_t_*, ***θ***) ∝ *p*(*y_t_*|*x_t_*, ***θ***)*p*(*x_t_*, ***θ***) and makes a single decision minimizing a cost function *u_t_* = argmin *∫ J*(*u_t_*, *x_t_*, ***θ***)*p*(*x_t_*|*y_t_*, ***θ***)*dx_t_*. In SDT, for example, observations are commonly assumed to be Gaussian distributed and the cost function *J* assigns a value to correct and wrong decisions. These assumptions make it straightforward to compute the observer’s decision probabilities *p*(*u_t_*|*x_t_*) given parameters ***θ*** (e.g. sensitivity and criterion in SDT) and invert the model to infer those parameters from behavior, yielding a posterior distribution *p*(***θ***|*u*_1:*T*_, *x*_1:*T*_), see Figure 1 C. Similarly, a psychometric curve can be interpreted as the decision probabilities of a Bayesian observer with a particular choice of measurement distribution, which determines the shape of the curve. The assumption of independence between trials is critical in the application of these modeling techniques.

**Figure 1.**
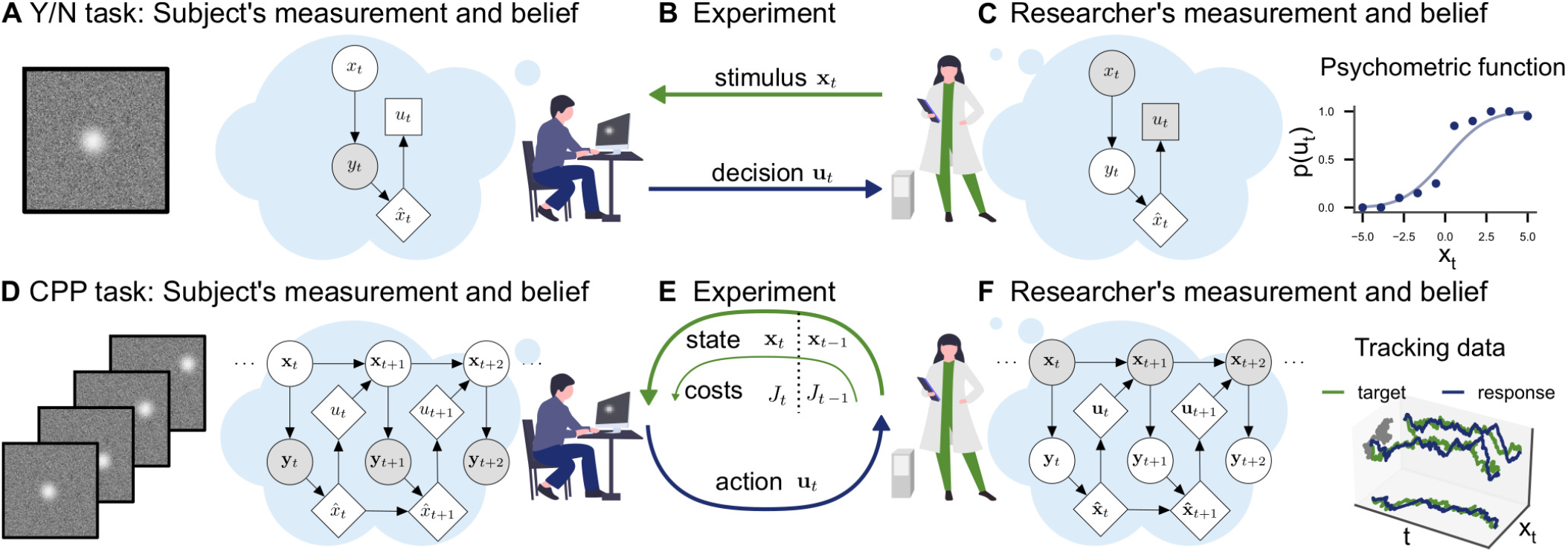
Conceptual frameworks for classical and continuous psychophysics. **A** In a classical psychophysics task, the subject receives stimuli *x_t_* on independent trials, generates sensory observations *y_t_*, forms beliefs about the stimulus 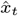 and **B** makes a single decision *u_t_* (e.g. whether the target stimulus was present or absent). **C** The researcher has a model of how the agent makes decisions and measures the subject’s sensitivity and decision criterion by inverting this model. For example, one can estimate the agent’s visual uncertainty by computing the width of a psychometric function. **D** In a continuous psychophysics task, a continuous stream of stimuli is presented. The subject has an internal model of the dynamics of the task, which they use to form a belief about the state of the world and then perform continuously actions based on their belief and subjective costs. **F** The researcher observes the subject’s behavior and inverts this internal model, e.g. using Bayesian inference applied to optimal control under uncertainty.

Continuous psychophysics abandons the independence of stimuli between individual trials. Instead, stimuli are presented in a continuous succession. For example, in a position tracking task, the stimulus moves from frame to frame and the observer’s goal is to track the stimulus with their mouse cursor (Figure 1 D, E). In a computational model of a continuous task, the stimulus dynamics are characterized by a state transition probability *p*(**x**_*t*_|**x**_*t*–1_, **u**_*t*–1_,***θ***) and can potentially be influenced by the agent’s actions **u**_*t*–1_. As in the trial-based task, the agent receives a stochastic measurement **y**_*t*_ ~ *p*(**y***_t_*|**x**_*t*_, ***θ***) and has the goal to perform a sequence of actions **u**_1:*T*_ = argmin *∫ J*(**u**_1:*T*_, **x**_1:*T*_)*p*(**x**_1:*T*_|**y**_1:*T*_)*d***x**_1:*T*_, which optimize a cost function. Formally, this problem of acting under uncertainty is a POMDP and is computationally intractable in the general case (Åström, 1965; Kaelbling et al., 1998).

In target tracking tasks used in previous continuous psychophysics studies (Bonnen et al., 2015; Bonnen et al., 2017; Huk et al., 2018; Knöll et al., 2018), in which the target is on a random walk, the dynamics of the stimulus are linear and subjects’ goal is to track the target. Tracking can be reasonably modeled with a quadratic cost function penalizing the separation between the target and the position of the tracking device. The variability in the dynamics and the uncertainty in the observation model can be modeled with Gaussian distributions. The resulting linear-quadratic-Gaussian (LQG) control problem (see Methods) is a well-studied special case of the general POMDP setting, which can be solved exactly by determining an optimal estimator and an optimal controller (B. D. Anderson & Moore, 2007; Tse, 1971). The optimal estimator is the KF and the optimal controller is the linear-quadratic regulator (LQR, Kalman, 1964).

We consider four models, each of which is a generalization of the previous one. Importantly, this means that the most general model contains the remaining three as special cases. First, the KF can be understood as the ideal observer in the tracking task (Figure 2 A): the agent’s response is simply the component of the estimate 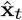, which corresponds to the expectation of the target’s position. The KF has only one parameter: perceptual uncertainty *σ*. Second, we use an optimal control model, which in addition to perceptual uncertainty about the target has perceptual uncertainty about the position of the cursor *σ_p_* and action variability *σ_m_* (Figure 2 B). Third, we use a bounded actor with internal behavioral costs *c* (Figure 2 C), which may penalize large actions and thus result in a larger lag between the response and the target. The control cost parameter *c* can implement a trade-off between tracking the target and expending effort (see Figure S1). Finally, we use a version of the bounded actor that bases its decisions on an internal model of the stimulus dynamics, which may differ from the true generative model employed by the experimenter (Figure 2 D). For example, in an experimental design with a target moving according to a random walk on position (Bonnen et al., 2015) with a true standard deviation *σ*_rw_, the subject could instead assume that the target follows a random walk on position with standard deviation *σ_s_* and an additional random walk on the velocity with standard deviation *σ_v_*. For a complete specification of the linear dynamical systems defining these models, see Table S1.

**Figure 2.**
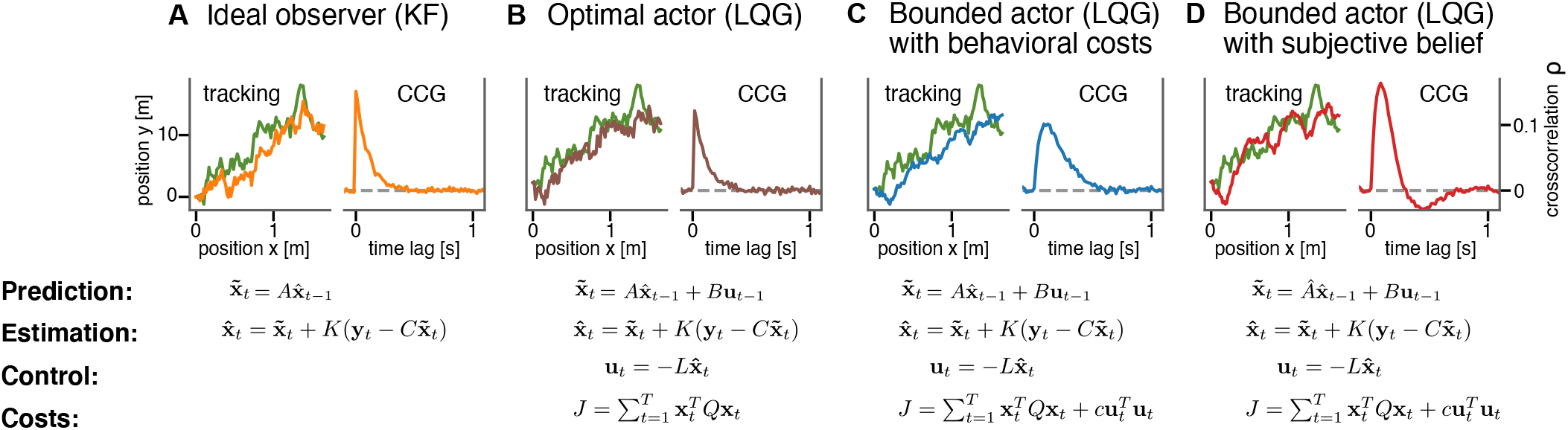
Computational models for continuous psychophysics. **A** In the KF model, the subject makes an observation **y**_*t*_ with Gaussian variability at each time step. They combine their prediction 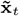 with their observation to compute an optimal estimate 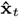. **B** In the optimal control models, this estimate is then used to compute an optimal action **u**_*t*_ using the LQR. The optimal action can be based on the task goal only or **C** bounded by internal costs, which e.g. penalize large movements. **D** Finally, the subject may act rationally using optimal estimation and control, but may use a subjective internal model of stimulus dynamics which differs from the true generative model of the task. These four different models are illustrated with an example stimulus and tracking trajectory (left subplots) and corresponding cross-correlograms (CCG, right subplots, see Appendix E).

### 2.2 Bayesian inverse optimal control

The normative models described in the previous section result in different predictions of how an optimal actor should behave in a continuous psychophysics task. From the point of view of a researcher, the goal is to estimate parameters ***θ***, which describe the perceptual, cognitive, and motor processes of the subject, given observed trajectories **x**_1:*T*_ (Figure 1 F) for each model. In the case of continuous psychophysics, these parameters include the perceptual uncertainty about the target *σ* and about one’s own cursor position *σ_p_*, the control cost *c*, and the action variability *σ_m_*, as well parameters which describe how the subject’s internal model differs from the true generative model. These properties are not directly observed and can only be inferred from the subject’s behavior.

To compute the posterior distribution according to Bayes theorem *p*(***θ***|**x**_1:*T*_) ∝ *p*(**x**_1:*T*_|***θ***)*p*(***θ***), we need the likelihood *p*(**x**_1:*T*_|***θ***), which we can derive using the Bayesian network model from the researcher’s point of view (Figure 1 F). The graph’s structure is identical to that describing the subject’s point of view. However, because different variables are observed, the decoupling of the perceptual and control processes no longer holds and we need to marginalize out the latent internal observations of the subject: *p*(**x**_1:*T*_|***θ***) = *∫*(**x**_1:*T*_, **y**_1:*T*_|***θ***)*d***y**_1:*T*_. Section 4.2 describes how to do this efficiently. We compute posterior distribution using probabilistic programming and Bayesian inference (see Section 4.3 A).

### 2.3 Simulation results

To establish that the inference algorithm can recover the model parameters from ground truth data, we generated 200 sets of parameters from uniform distributions resulting in realistic tracking data (see Appendix F.1 for details). For each set of parameters, we simulated data sets consisting of 10, 25, 50, and 100 trials (corresponding to 2, 5, 10, and 20 minutes) from the bounded actor model. Figure 3 A shows one example posterior distribution. Average posterior means and credible intervals relative to the true value of the parameter are shown in Figure 3 B. With only 2 minutes of tracking, the 95% posterior credible intervals of the perceptual uncertainty *σ* are [0.8, 1.3] relative to the true value on average. This means, that the tracking data of an experiment lasting 2 minutes is sufficient to obtain posterior distributions over model parameters for which 95% of probability are within a range of 20% underestimation and 30% overestimation of the true values. As a comparison, we consider the simulations for estimating the width of a psychometric function in classical psychophysical tasks conducted by Schütt et al. (2015). With 800 forced-choice trials (corresponding to roughly 20 minutes), the 95% posterior credible intervals are within [0.6, 1.6] relative to the true value on average. This suggests a temporal gain for continuous psychophysics of at least a factor of 10. These simulation results are in accordance with the empirical results of Bonnen et al. (2015), who had shown that the tracking task yields stable but biased estimates of perceptual uncertainty in under two minutes of data collection.

**Figure 3.**
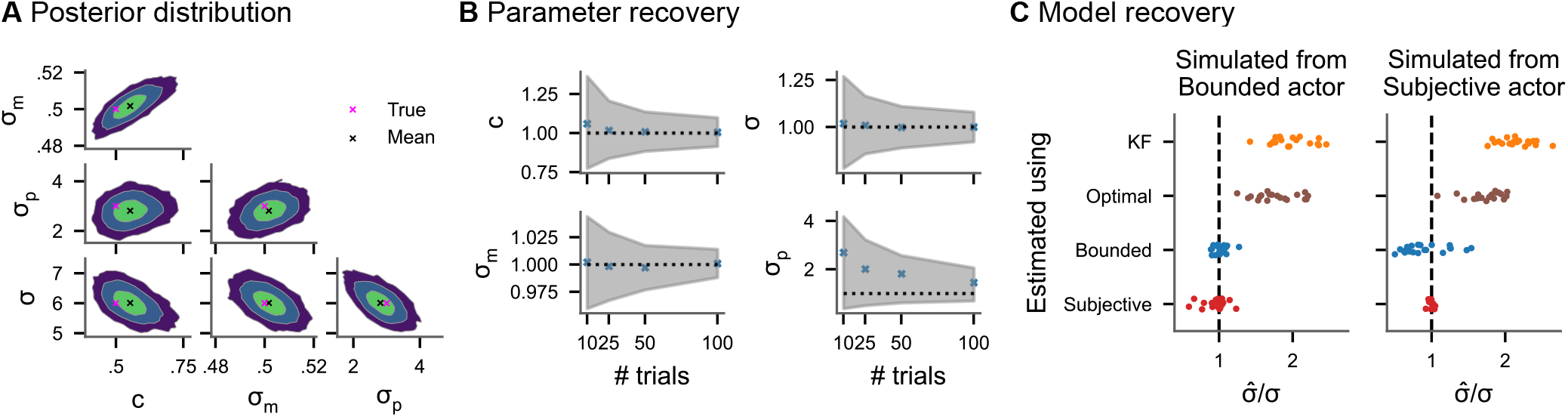
Inference on simulated data. **A** Pairwise joint posterior distributions (0.5, 0.9 and 0.99 highest density intervals) inferred from simulated data with parameters representative of real tracking data (*c* = 0.5, *σ* = 10.0, *σ_p_* = 8.0 and *σ_m_* = 0.5). The green dots mark the true value used to simulate the data, while the black lines mark the posterior mean. **B** Average posterior means and average 95% credible intervals relative to the true value for different numbers of trials (200 repetitions each). **C** Model recovery analysis. Inferring perceptual uncertainty (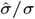, posterior mean relative to the true value) with each of the four models from data simulated from the bounded actor model and subjective actor model, respectively.

Behavior in a task with an interplay of perception and action such as a tracking task contains additional sources of variability and bias beyond sensory influences: repeated actions with the same goal are variable and the cost of actions may influence behavior. We model these factors as action variability *σ_m_* and control cost *c*, respectively, in the bounded actor model. A model without these factors needs to attribute all the experimentally measured behavioral biases and variability to perceptual factors, even when they are potentially caused by additional cognitive and motor processes. We substantiate this theoretical argument via simulations. We simulated 20 datasets with different values for *c* and *σ_m_* sampled from uniform ranges from the bounded actor model and the subjective model. Then, we fit each of the four different models presented in Figure 2 to the tracking data and computed posterior mean perceptual uncertainties. The results are shown in Figure 3 C. The posterior means of *σ* from the bounded actor and subjective model scatter around the true value, with the model used to simulate the data being the most accurate for estimating *σ*. The other two models (KF and optimal actor), which do not contain intrinsic behavioral costs, both overestimate *σ*. This overestimation is worse when the model does also not account for action variability (KF). Furthermore, Bayesian model comparison confirms that the model used to simulate the data is also the model with the highest predictive accuracy in all cases (Figure S2).

### 2.4 Continuous psychophysics

We now show that the ability of the optimal control models to account for control costs and action variability results in more accurate estimates of perceptual uncertainty in a target-tracking task. We reanalyze data from three subjects in two previously published experiments (Bonnen et al., 2015, see Appendix G.1 for details): a position discrimination and a position tracking task. Both experiments employed the same visual stimuli: Gaussian blobs with six different widths within white noise backgrounds were used to manipulate the stimulus contrast (see Figure 4 A). In the 2IFC discrimination task, the psychophysical measurements of perceptual uncertainty increased with decreasing contrast (Figure 4 A, green lines). In the tracking experiment, the same visual stimuli moved on a Gaussian random walk with a fixed standard deviation and subjects tracked the target with a small red mouse cursor.

**Figure 4.**
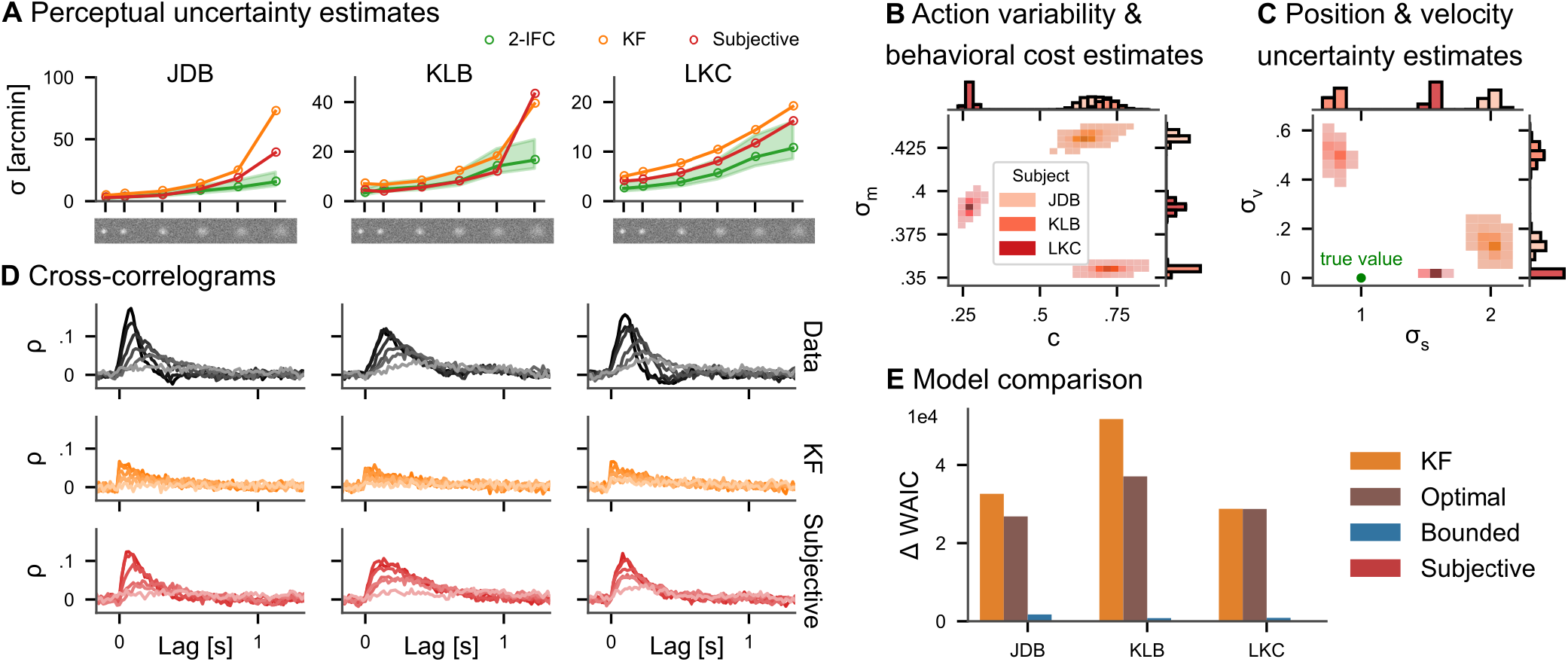
Continuous psychophysics. **A** Perceptual uncertainty (*σ*) parameter estimates (posterior means) for the 2IFC task and the tracking task (KF and LQG models) in the six blob width conditions (Bonnen et al., 2015). The shaded area represents the range of observed deviations between 2IFC and YN tasks in the literature (Yeshurun et al., 2008). **B** Posterior distributions for action cost (*c*) and action variability (*σ_m_*). **C** Posterior distributions for subjective stimulus dynamics parameters (position: *σ_s_*, velocity: *σ_v_*). The true values of the target’s random walk are marked by dashed lines. **D** CCGs of the empirical data and both models for all three subjects. **E** Model comparison. The difference in WAIC w.r.t. the best model is shown with error bars representing WAIC standard error. Models without control cost (KF and optimal) fare worst, while the subjective model has the highest predictive accuracy.

We fit all four models presented in Figure 2 to the data of the tracking experiment. To a first approximation, it is reasonable to assume that action variability, perceptual uncertainty about the mouse position, and control costs are independent of the target blob width, so that these parameters are shared across the different contrast conditions. The perceptual uncertainty *σ* should be different across contrasts, so we inferred individual parameters *σ* per condition. We assume ideal temporal integration over individual frames, including for the reanalysis of the KF model (see Appendix D.1), which certainly constitutes a lower bound on modeled perceptual uncertainty. We focus on the KF and the subjective version of the LQG model in the following.

The posterior means of estimated perceptual uncertainty in both models were highly correlated with the classical psychophysical measurements (r > 0.88 for both models and all subjects). However, the parameters from the LQG model are smaller in magnitude than KF’s in all three subjects and all but one blob width condition (Figure 4 A). The reason for this is that the KF attributes all variability to perceptual uncertainty, while the other models explicitly include cognitive and motor influences (see Section 2.3). The average factor between the posterior mean perceptual uncertainty in the continuous task and the 2IFC task in the 5 higher contrast conditions is 1.20 for the LQG model, while it is 1.73 for the KF. Only in the lowest contrast condition, it increases to 2.20 and 2.93 arcmin, respectively. Thus, accounting for action variability and control effort in our model leads to estimates of perceptual uncertainty, which are closer to those obtained with the classical 2IFC psychophysics task. Note that a deviation between the two psychophysical tasks should not be surprising based on previous research comparing the thresholds obtained with different traditional paradigms, as we further discuss below.

Importantly, in addition to the perceptual uncertainty parameter, we also obtain posterior distributions over the other model parameters (Figure 4 B). The three subjects differ in how variable their actions are (*σ_m_*) and how much they subjectively penalize control effort (*c*). Additionally, the bounded actor model with subjective beliefs about the stimulus dynamics infers subjects’ implicit assumptions about how the target blob moved. The model estimates that two of the three subjects assume a velocity component *σ_v_* to the target’s dynamics, although the target’s true motion according to the experimenter’s generative model does not contain such a motion component, while in the third participants this estimated velocity component is close to zero. Similarly, the model also infers the subjectively perceived randomness of the random walk, i.e. the standard deviation *σ_s_* (Figure 4 C), for each subject, which can be smaller or larger than the true standard deviation used in generating the stimulus.

As an important test, previous research has proposed comparing the autocorrelation structure of the tracking data with that of the model’s prediction (Bonnen et al., 2015; Huk et al., 2018; Knöll et al., 2018). We simulated data from both the KF model and the subjective model (20 trials, using the posterior mean parameter estimates) and compared it to the empirical data. To this end, we computed cross-correlograms (CCGs, i.e. the cross-correlation between the velocities of the target and the response, see Appendix E) for each trial of the real data and simulated data from both models (Figure 4 C). Indeed, the LQG models capture more of the autocorrelation structure of the tracking data, quantified by the correlation between the CCGs of the model and those of the data, which was 0.61 for the KF and 0.86 for the subjective actor. To quantitatively compare the models, we employ Bayesian model comparison using the widely applicable information criterion (WAIC; Watanabe & Opper, 2010, see Section 4.4). WAIC is computed from posterior samples and estimates the out-of-sample predictive accuracy of a model thereby taking into account that models with more parameters are more expressive. Overwhelming evidence for the subjective bounded actor model is provided by Akaike weights larger than 0.98 in all three subjects, with the weight for the KF model being equal to 0 within floating point precision. This means that the behavioral data was overwhelmingly more likely under the bounded actor model with subjective stimulus beliefs than the KF model, even after taking the wider expressivity of the model into account. Such strong evidence in favor of one of the models is the consequence of the large number of data samples obtained in continuous psychophysics paradigms.

### 2.5 Application to other datasets

We furthermore applied all four models to data from an additional experiment in which subjects tracked a target with their finger using a motion tracking device (Bonnen et al., 2017, see Appendix G.2 for details). Comparisons of CCGs for one subject are shown in Figure 5 A (see Figure S3 for all subjects). Qualitatively, the subjective model captures the cross-correlation structure of the data better than the other two models. To quantitatively evaluate the model fits, we again used WAIC (Figure 5 B). In two of the three subjects, the subjective model performs best, while in one subject the bounded actor with accurate stimulus model accounts better for the data. Cross-validation provides additional support for the subjective model, since it gives more consistent estimates of perceptual uncertainty (Figure S4). Again, we can inspect and interpret the other model parameter such as control costs, action variability and subjective belief about target dynamics (Figure S5). Similarly, when applied to another experiment, in which one human participant, two monkeys, and a marmoset tracked optic flow fields with their gaze (Knöll et al., 2018), model selection favored the bounded actor with subjective beliefs about stimulus dynamics (Figure S6).

**Figure 5.**
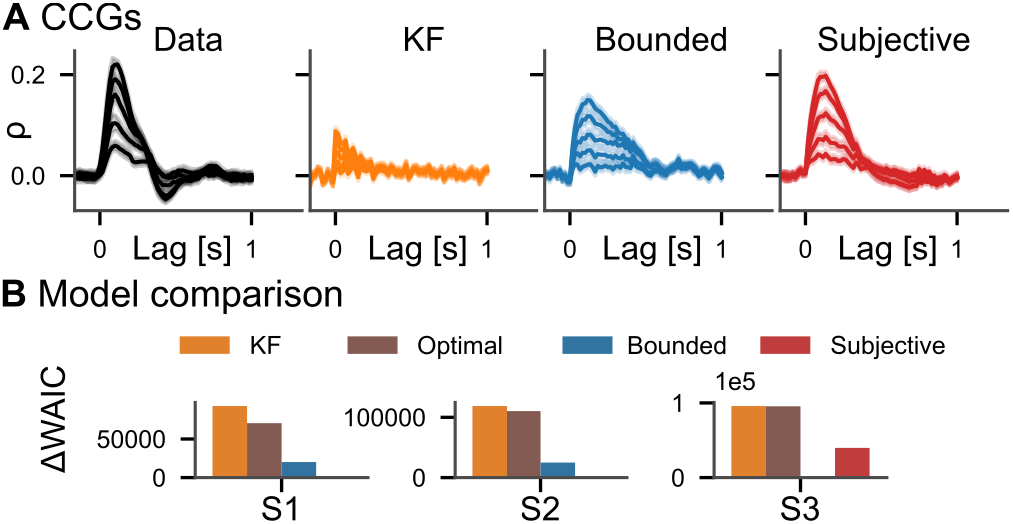
Model comparison on motion tracking data. **A** Average CCGs for the S1 from Bonnen et al. (2017) and three models in different target random walk conditions. For the other two subjects, see Fig. S3. **B** Difference in WAIC w.r.t. best model.

## 3 Discussion

We introduce an analysis method for continuous psychophysics experiments (Bonnen et al., 2015; Bonnen et al., 2017; Huk et al., 2018; Knöll et al., 2018), which allows estimating subjects’ perceptual uncertainty, the quantity of interest in the analysis of traditional psychophysics experiments with SDT. We validated the method on synthetic data as well as on experimental data, which were collected together with equivalent stimuli in a 2IFC paradigm for the purpose of empirical validation (Bonnen et al., 2015). While perceptual uncertainties recovered with previous analysis methods were larger by one order of magnitude compared to those recovered with SDT (Bonnen et al., 2015), we infer values that differ by a factor of 1.36 on average. This deviation is well within the bounds of differences between classical psychophysical methods, where Yeshurun et al. (2008) have e.g. reported deviations between 0.8 and 1.5 between 2AFC and Yes-No tasks (cf. Appendix A). The feasibility of the present analysis method was demonstrated for actions involving participants’ tracking with a computer mouse (Bonnen et al., 2017), tracking by pointing a finger (Bonnen et al., 2015), and tracking using gaze (Knöll et al., 2018).

Because subjects’ behavior is conceptualized as optimal control under uncertainty (B. D. Anderson & Moore, 2007; Åström, 1965; Hoppe & Rothkopf, 2019; Kaelbling et al., 1998), the optimal actor model additionally contains action variability and a cost function encapsulating the behavioral goal of tracking the target. The present analysis method probabilistically inverts this model similarly to approaches in inverse reinforcement learning (Ng, Russell, et al., 2000; Rothkopf & Dimitrakakis, 2011; Ziebart et al., 2008) and inverse optimal control (Chen & Ziebart, 2015; Herman et al., 2016; Schmitt et al., 2017). The inferred action variability can be attributed to motor variability (Faisal et al., 2008) but also to other cognitive sources, including decision variability (Gold & Shadlen, 2007). We extended this model to a bounded actor model in the spirit of rational analysis (J. R. Anderson, 1991; Gershman et al., 2015; Simon, 1955) by including subjective costs for carrying out tracking actions. These behavioral costs correspond to intrinsic, subjective effort costs, which may include biomechanical costs (Di Prampero, 1981) as well as cognitive effort (Shenhav et al., 2017), trading off with the subject’s behavioral goal of tracking the target.

The model was furthermore extended to allow for the possibility that subjects may act upon beliefs about the dynamics of the target stimuli that differ from the true dynamics employed by the experimenter. Such a situation may arise due to perceptual biases in the observation of the target’s dynamics. Model selection using WAIC (Watanabe & Opper, 2010) favored this bounded actor with subjective beliefs about stimulus dynamics in all four experiments involving human and monkey subjects with a single exception, in which the bounded actor model better accounted for the behavioral data. Thus, we found overwhelming evidence that subjects’ behavior in the continuous psychophysics paradigm was better explained by a model that includes action variability and internal costs for actions increasing with the magnitude of the response.

One possible criticism of continuous psychophysics is that it introduces additional unmeasured factors such as action variability, intrinsic costs, and subjective internal models. While classical psychophysical paradigms take great care to minimize the influence of these factors by careful experimental design, they can nevertheless still be present, e.g. as serial dependencies (Fischer & Whitney, 2014; Fründ et al., 2014; Green, 1964). Similarly, estimates of perceptual uncertainty often differ between classical psychophysical tasks when compared directly. Besides the difference between 2AFC and Yes-No tasks mentioned above, there are differences between 2AFC tasks, in which two stimuli are separated spatially, and their equivalent two-interval forced-choice (2IFC) tasks, in which the stimuli are separated in time (Jäkel & Wichmann, 2006). While the former task engages spatial attention, the latter engages memory. These factors are typically also not accounted for in SDT-based models. On a conceptual level, this underscores the fact that psychophysical methods always imply a generative model of behavior (Green & Swets, 1966; Swets, 1986; Wixted, 2020), in which quantities besides sensitivity and bias can play a role.

The limitations of the current method are mainly due to the limitations of the LQG framework. For instance, the assumption that perceptual uncertainty is constant across the whole range of the stimulus domain is adequate for position stimuli, but would not be correct for stimuli which behave according to Weber’s law (Weber, 1834). This could be addressed using an extension of LQG to signal-dependent variability (Todorov, 2005) to which our inverse optimal control method can be adapted (Schultheis et al., 2021). Similarly, the assumption of linear dynamics can be overcome using generalizations of the LQG framework (Todorov & Li, 2005) or policies parameterized with neural networks (Kwon et al., 2020).

Taken together, the current analysis framework opens up the prospect of a wider adoption of continuous psychophysics paradigms in psychology, cognitive science, and neuroscience, as it alleviates the necessity of hundreds of repetitive trials with binary forced-choice responses in expert observers, recovering perceptual uncertainties well within the bounds of classic paradigms and SDT. Additionally, it extracts meaningful psychological quantities capturing behavioral variability, effort costs, and subjective perceptual beliefs and provides further evidence for the importance of modeling the behavioral goal and subjective costs in experiments seemingly only investigating perception, as perception and action are inseparably intertwined.

## 4 Methods

### 4.1 Linear-quadratic-Gaussian control problem

The LQG control problem (B. D. Anderson & Moore, 2007) is defined by a discrete-time linear dynamic system with Gaussian noise, a linear observation model with Gaussian noise, and a quadratic cost function. The equation linear dynamical system with control inputs is

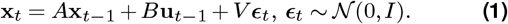

The linear observation model is

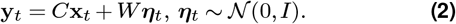

The quadratic cost function is

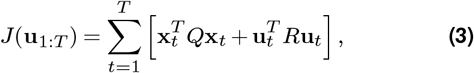

where *Q* defines the costs for the state (e.g. squared distance between target and mouse / hand) and *R* defines the cost of actions (biomechanical, cognitive etc.).

For the LQG control problem, the *separation principle* between estimation and control holds (Davis & Vinter, 1985). This means that the optimal solutions for the estimator (KF) and controller (LQR) can be computed independently from one another. The KF (Kalman, 1960) is used to form a belief 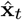 about the current state based on the previous estimate and the current observation

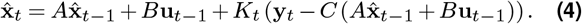

This belief is then used to compute the optimal action **u**_*t*_ based on the optimal control law of the LQR: 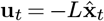. For the derivations of *K* and *L*, see Appendix B.

### 4.2 Likelihood function

Due to the conditional independence assumption implied by the graphical model from the researcher’s point of view (Figure 1 F), the next state **x**_*t*+1_ depends on the whole history of the subject’s noisy observations **y**_1:*t*_ and internal estimates 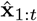. Since, for the researcher, the subject’s observations **y**_*t*_ are latent variables, the Markov property between the **x**_*t*_ no longer holds and each **x**_*t*_ depends on all previous **x**_1:*t*–1_, i.e. 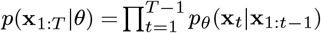. We can, however, find a more manageable representation of the dynamical system by describing how the **x**_*t*_ and 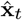 evolve jointly (Van Den Berg et al., 2011). This allows us to write down a joint dynamical system of **x**_*t*_ and 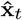, which only depends on the previous time step:

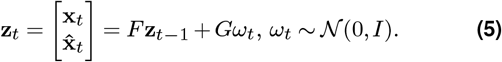

For the derivation of this joint dynamical system and the definitions of *F* and *G*, see Appendix C.1. The derivation is extended to account for differences between the true generative model and the agent’s subjective internal model in Appendix C.2.

This allows us to recursively compute the likelihood function in the following way. If we know the distribution 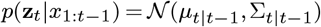, we can compute the joint distribution *p*(**z**_*t*_,**z**_*t*+1_|**x**_1:*t*–1_) using the dynamical system from Eq. (5). Since this distribution is a multivariate Gaussian, we can condition on **x**_*t*_ and marginalize out the 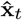 and 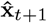 to arrive at *p_θ_*(**x**_*t*+1_|**x**_1:*t*_). For details, see Appendix C. The computation of the likelihood function involves looping over a possibly large number of time steps. To make computing gradients w.r.t. the parameters computationally tractable, we use the Python automatic differentiation library `jax` (Frostig et al., 2018). Our implementation will be made available on github.

### 4.3 Bayesian inference

Equipped with the likelihood function derived above, we can use Bayes’ theorem to compute the posterior distribution for the parameters of interest:

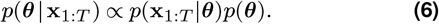

Since most parameters (except the internal costs *c*) are interpretable in degrees of visual angle, prior distributions *p*(***θ***) can be chosen in a reasonable range depending on the experimental setup. We used half-Cauchy priors with the scale parameter *γ* set to 50 for *σ*, 25 for *σ_p_*, 0.5 for *σ_m_* and 1 for *c*. We verified that the prior distributions lead to reasonable tracking data using prior predictive checks with the cross-correlograms of the tracking data.

Samples from the posterior are drawn using NUTS (Hoffman & Gelman, 2014) as implemented in the probabilistic programming package numpyro (Phan et al., 2019).

### 4.4 Model comparison

We compare models using the widely-applicable information criterion (WAIC; Watanabe & Opper, 2010), which approximates the expected log pointwise predictive density. Importantly, unlike AIC it makes not parametric assumptions about the form of the posterior distribution. It can be computed using samples from the posterior distribution (see Vehtari et al., 2017, for a detailed explanation):

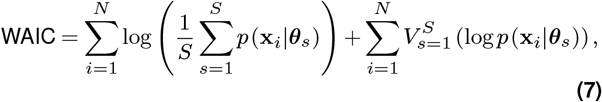

where 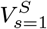 is the variance over samples from the posterior. Since WAIC requires individual data points **x**_*i*_ to be independent given the parameters ***θ***, we define the terms *p*(**x**_*i*_|***θ***_*s*_) as the likelihood of a trial *i* given posterior sample *s*, which is computed as explained in Section 4.2. We use the implementation from the Python package `arviz` (Kumar et al., 2019).

When comparing multiple models, the WAIC values can be turned into Akaike weights summing to 1

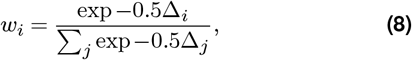

where Δ_*i*_ is the difference between the current model’s and the best model’s WAIC. The Akaike weights can be interpreted as the relative likelihood of each model.

## Acknowledgments

We thank Kathryn Bonnen and Lawrence Cormack for providing their behavioral data. Calculations for this research were conducted on the Lichtenberg high performance computer of the TU Darmstadt. This research was supported by “The Adaptive Mind”, funded by the Excellence Program of the Hessian Ministry of Higher Education, Science, Research and Art.

## Supplementary information

## A Differences in measured sensitivities across psychophysical paradigms

Yeshurun et al. (2008) reviewed a large body of psychophysical studies involving analyses with Signal Detection Theory (SDT). Specifically, they investigated the validity of the theoretical result (Green & Swets, 1966), that the forced-choice sensitivity 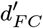 differs from the sensitivity obtained through an equivalent Yes-No task 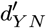 by a factor of 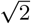, i.e. 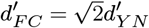. Apart from running a series of experiments targeted at evaluating the validity of this theoretical result, the authors reported the actual values of this factor based on an extensive literature review involving numerous studies investigating perceptual sensitivities across sensory modalities. Indeed, the literature review collected a broad range of deviations from the theoretical value. Of particular interest are here those studies, which provide an upper and lower bound on the empirically found sensitivities differences between these two paradigms.

The authors report a study by Jesteadt and Bilger (1974), which investigated auditory frequency discrimination and intensity discrimination with both 2-IFC and Yes–No tasks. This study found that both frequency and intensity discrimination performance in the 2-IFC task was better than performance in the equivalent Yes–No task, albeit not by a factor of approximately 1.41 but by a factor of 2.13, corresponding to a 1.5 fold deviation between the results obtained in the two paradigms relative to the theoretical prediction.

On the other end of the spectrum, the authors report a study by Markowitz and Swets (1967), which compared performance in auditory detection with both a 2-IFC and a Yes–No task. In this particular case, the empirical factor also deviated significantly from approximately 1.41 and was instead 1.15, corresponding to a 0.8 fold deviation between the results obtained in the two paradigms relative to the theoretical prediction.

## B LQG control derivations

For the LQG control problem as defined in Section 4.1, the *separation principle* between estimation and control holds (Davis & Vinter, 1985). This means that the optimal solutions for the estimator and controller can be computed independent from one another. The optimal estimator is the KF (Kalman, 1960)

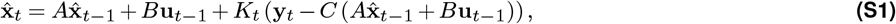

where the Kalman gain *K_t_* = *P_t_C^T^* (*CP_t_C^T^* + *WW^T^*)^−1^ is computed using a discrete-time Riccati equation

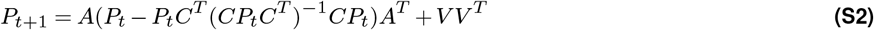

initialized with *P*_0_ = *VV^T^*. The optimal controller

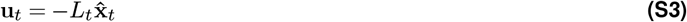

is the linear-quadratic regulator (LQR) *L_t_* = (*B^T^ S*_*t*+1_*B* + *R*)^−1^ *B^T^ S*_*t*+1_*A* and is also computed with a discrete-time Riccati equation

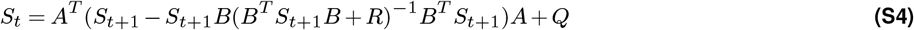

initialized with *S_T_* = *Q*. In principle, all matrices of the dynamical system and thus also *K* and *L* could be time-dependent, but since this is not the case in the models considered in this work, we leave the time-indices out for notational simplicity. Due to this time-invariance of the dynamical system, the gain matrices *K_t_* and *L_t_* typically converge after a few time steps. This allows us to use the converged versions, which we call *K* and *L*, in our inference method.

## C Inverse LQG likelihood

### C.1 Derivation

We define ***θ*** to be the vector containing any parameters of interest, which influence the LQG system defined above. This could be parameters of the matrices of the dynamical system (*A*, *B*, *V*), the observation model (*C*, *W*), or the cost function (*Q*, *R*). For notational simplicity, we do not explicitly indicate this dependence of the matrices on the parameters. We are interested in the posterior probability

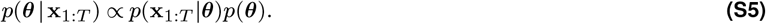

We start by writing the likelihood using repeated applications of the chain rule:

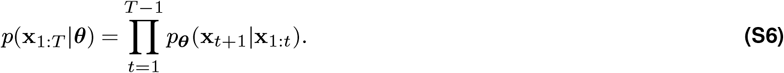

The current state **x**_*t*_ depends on the whole history of previous states **x**_1:*t*–1_, since the internal observations **y**_1:*T*_ and estimates 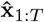 are unobserved from the researcher’s perspective (see Figure 1 F). The linear Gaussian assumption, however, allows us to marginalize over these variables efficiently. To do this, we start by expressing the current state **x**_*t*_ and estimate 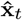 such that they only depend on the previous time step. This idea is based on LQG-MP (Van Den Berg et al., 2011), a method for planning in LQG problems, which computes the distribution of states and estimates without any observations. We start by inserting the LQR control law (Eq. (S3)) into the state update (Eq. (1)):

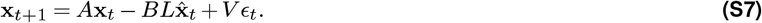

Then, we rewrite the KF update equation (Eq. (S1))

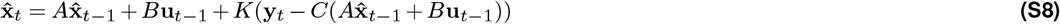

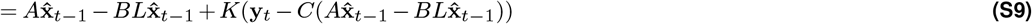

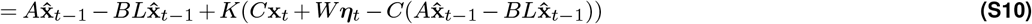

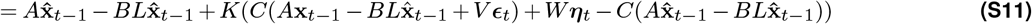

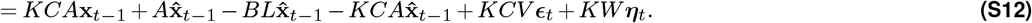

Here, we have again inserted the LQR control law (Eq. (S3)), then the observation model (Eq. (2)), then the state transition model (Eq. (1)) and finally rearranged the terms. Putting these together, we can write

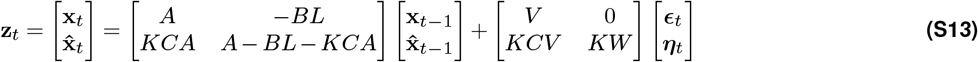

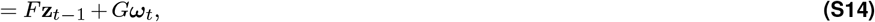

This allows us to recursively compute the likelihood function (Eq. (S6)) in the following way. If we know the distribution 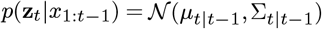, we can compute the joint distribution *p*(**z**_*t*_, **z**_*t*+1_|**x**_1:*t*–1_) using the dynamical system from Eq. (S14) and the formulas for linear transformations of Gaussian distributions, giving us the mean and covariance matrix

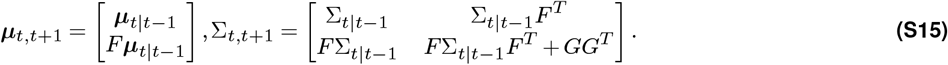

Since this distribution is a multivariate Gaussian, we can condition on **x**_*t*_ to obtain 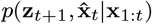 and marginalize out 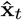 to arrive at *p*_***θ***_(**z**_*t*+1_|**x**_1:*t*_). We do this by partitioning the covariance matrix as

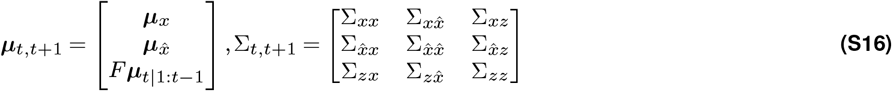

and applying the equations for conditioning in a multivariate Gaussian distribution to obtain

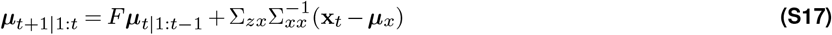

and

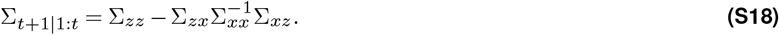

These ***μ***_*t*+1|1:*t*_ and ∑_*t*+1|1:*t*_ are then used for *p*(**z**_*t*+1_|*x*_1:*t*_) in the next iteration. The contribution to the likelihood at the current time step *p*(**x**_*t*+1_|**x**_1:*t*_) is obtained by marginalizing this distribution over 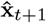. Without loss of generality (since we can subtract the initial time step from the observed trajectories to let them start at zero), we initialize *p*(**z**_1_|**x**_0_) with ***μ***_1|0_ = 0 and ∑_1|0_ = *GG^T^*.

### C.2 Extension to subjective internal models

If we allow the subjective internal model to differ from the true generative model, we need to adapt the model equations accordingly. We denote matrices of the true generative model with superscript *g* and matrices representing the subjective internal model with superscript *s*. The state update model

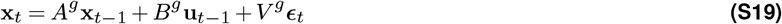

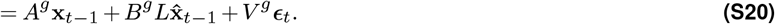

and the observation model

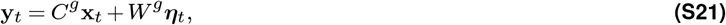

remain the same and contain only the matrices of the generative model. The KF update equation, being internal to the subject, is now influenced by several subjective components

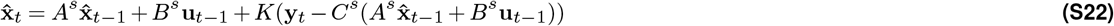

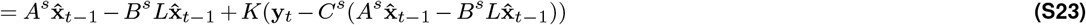

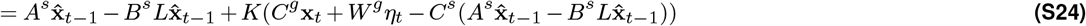

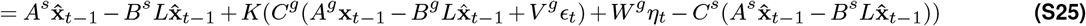

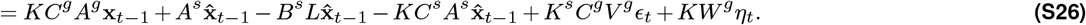

The joint dynamical system of **x**_*t*_ and 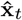 is then

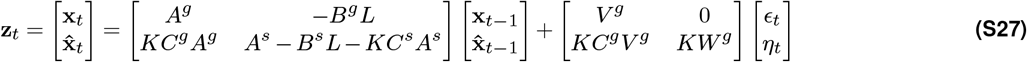

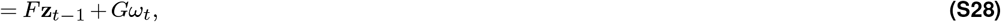

Then, we proceed by conditioning on the observed states and marginalizing out the subject’s internal estimates at each time step as described in the previous subsection.

## D Models of the tracking task

As explained in Section 2.1, we consider a succession of increasingly more complicated models, which can be seen as generalizations of the previous ones. In this section, we provide a verbal description of each of the models. Table S1 contains full definitions of all the matrices needed to define the models.

**Table S1.** Model overview

| Model | Dynamical system matrices | Cost function | Free parameters |
| --- | --- | --- | --- |
| Ideal observer | $A = \begin{bmatrix} 1 \end{bmatrix}, B = \begin{bmatrix} 0 \end{bmatrix}, V = \begin{bmatrix} \sigma_{rw} \end{bmatrix},$<br>$C = \begin{bmatrix} 1 \end{bmatrix}, W = \begin{bmatrix} \sigma \end{bmatrix}$ | - | $\sigma$ |
| Optimal actor | $A = \begin{bmatrix} 1 & 0 \\ 0 & 1 \end{bmatrix}, B = \begin{bmatrix} 0 \\ dt \end{bmatrix}, V = \begin{bmatrix} \sigma_{rw} & 0 \\ 0 & \sigma_m \end{bmatrix}$<br>$C = \begin{bmatrix} 1 & 0 \\ 0 & 1 \end{bmatrix}, W = \begin{bmatrix} \sigma & 0 \\ 0 & \sigma_p \end{bmatrix}$ | $Q = \begin{bmatrix} 1 & -1 \\ -1 & 1 \end{bmatrix}, R = \begin{bmatrix} 0 \end{bmatrix}$ | $\sigma, \sigma_m, \sigma_p$ |
| Bounded actor | $A, B, V, C, W$ as above | $Q$ as above, $R = \begin{bmatrix} c \end{bmatrix}$ | $\sigma, \sigma_m, \sigma_p, c$ |
| Subjective actor | $A, B, V, C, W$ as above<br>$A^s = \begin{bmatrix} 1 & 0 & dt \\ 0 & 1 & 0 \\ 0 & 0 & 1 \end{bmatrix}, B^s = \begin{bmatrix} 0 \\ dt \\ 0 \end{bmatrix}, V^s = \begin{bmatrix} \sigma_s & 0 & 0 \\ 0 & \sigma_m & 0 \\ 0 & 0 & \sigma_v \end{bmatrix}$<br>$C^s = \begin{bmatrix} 1 & 0 & 0 \\ 0 & 1 & 0 \end{bmatrix}, W^s = W$ | $Q, R$ as above<br>$Q^s = \begin{bmatrix} 1 & -1 & 0 \\ -1 & 1 & 0 \\ 0 & 0 & 0 \end{bmatrix},$<br>$R^s = R$ | $\sigma, \sigma_m, \sigma_p, c, \sigma_s, \sigma_v$ |

### D.1 Ideal observer (KF)

For the KF model, the state is simply the position of the target, 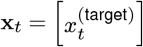. The subject’s cursor position is taken as the internal estimate of the state 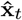. The likelihood of the observed data 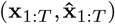 can be computed as above (Appendix C), but since the model does not explicitly include actionsand instead treats the internal estimate is as the subject’s cursor position, we need to condition on **x**_1:*T*_ and 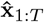) at every time step.

### D.2 Optimal and bounded actor

The models based on LQG control explicitly include the subject’s cursor position as a part of the state. In the simplest case, the state is two-dimensional and contains the position of the target stimulus and the subject’s position of the response: 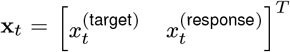. The continuous psychophysics experiments considered here used target stimuli moving according to a simple Gaussian random walk. Accordingly, the model may assume that the target only moves because of the random walk with standard deviation *σ*_rw_ and that the response is influenced by the agent’s control input and by action variability *σ_m_*. The agent observes the target and response position separately, with independent observation noise in both dimensions (*σ* for the target and *σ_p_* for the response). The behavioral cost of actions is penalized by the parameter *c*. For the optimal actor without internal costs, we set *c* = 0, while for the bounded actor *c* is a free parameter.

### D.3 Subjective model

In the subjective model, the actual matrices of the generative model remain identical to the basic model, but the agent assumes a different dynamical system, in which the target has an additional dimension representing its velocity 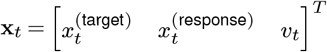. Furthermore, the subject may assume that in addition to the target position’s random walk, the target’s velocity changes over time according to its own random walk. The agent still receives an observation of the positions only, with independent observation noise on both dimensions. The parameters *σ_s_* and *σ_v_* represent the subject’s belief about the standard deviation of the target’s position and velocity.

## E Cross-correlograms

As a summary statistic of the tracking data, we use cross-correlograms as defined by Mulligan et al. (2013). The cross-correlogram at time lag *τ* is the correlation between the velocities of the target 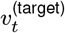 at time *t* and the velocity of the response 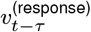 at time (*t* – *τ*):

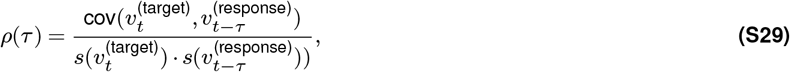

where cov(*x*, *y*) and *s*(*x*) are the covariance and standard deviation across all time steps and trials. The velocities are estimated from position data via finite differences.

## F Simulations

### F.1 Parameter recovery

To evaluate the inference method, 200 sets of parameters for the bounded actor model were sampled from the following uniform distributions

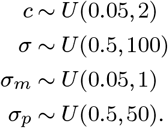

For each set of parameters, we simulated data sets with the following number of trials containing 10, 25, 50 and 100 trials. Each trial was 720 time steps long. For each data set, 4 Markov chains with 7500 samples each were drawn from the posterior distribution, after 2500 warm-up steps.

### F.2 Model recovery

To investigate how different models behave when fit to tracking data generated from one of the other models, we performed a model recovery analysis. To this end, 20 different values for the parameters *c* and *σ_m_* were sampled from uniform distributions defined above. The values the other model parameter were fixed at *σ* = 6, *σ_p_* = 1, *σ_s_* = 0.1 and *σ_v_* = 10. For each set of parameters, we simulated datasets of 50 trials with 500 time steps per trial from the bounded actor model and the subjective model. Then, we estimated fit each of the four different models presented in Figure 2 to the tracking data. We ran 4 chains with 5000 samples each after 2000 warm-up steps.

## G Experiments

We reanalyze data from three previous publications (Bonnen et al., 2015; Bonnen et al., 2017; Knöll et al., 2018). In this section, we describe our data analysis procedures for these datasets. For detailed information about the experiments, we refer to the original publications.

### G.1 Continuous psychophysics

First, we reanalyze data from three subjects in two previously published experiments (Bonnen et al., 2015): a position discrimination and a position tracking task. The stimuli were “Gaussian blobs”, two-dimensional Gaussian functions on Gaussian pixel noise, which changed from frame to frame. The visibility of the stimuli was manipulated via the standard deviation of the Gaussian blobs, with larger standard deviations leading to lower luminance increments and thus to a weaker signal. The standard deviations were 11, 13, 17, 21, 25, and 29 arcmin.

In the discrimination experiment, position discrimination thresholds for each blob width were measured in a two-interval forced-choice (2IFC) task. We fit cumulative Gaussian psychometric functions with lapse rates in a Beta-Binomial model to the discrimination performance using `psignifit` (Schütt et al., 2015). We use the posterior mean of the standard deviation of the Gaussian psychometric function as our estimate of the perceptual uncertainty in the 2IFC task.

In the tracking experiment, the same visual stimuli moved on a random walk with a standard deviation of one pixel (1.32 arcmin of visual angle) per frame at a frame rate of 60Hz. Subjects were instructed to track the target with a small red mouse cursor. Each subject completed 20 trials for each blob width, with each trial lasting 1200 time steps (20 seconds). Each subject completed 20 trials for each blob width, with each trial lasting 1200 time steps (20 seconds). As in Bonnen et al. (2015), the response time series were shifted by *τ* = 12 time steps w.r.t. the target time series to account for a constant time lag. Instead, one could explicitly account for the time lag in the observation function of the model by extending the state space to include the *τ* previous time steps, but we found no appreciable differences in the model fits. We obtain posterior mean perceptual uncertainty estimates using the methods described above (Section 4.2, 4.3). Since there are two stimulus presentations in a 2IFC task and we are interested in perceptual uncertainty for a single stimulus presentation, we divided the estimates of the perceptual uncertainty by 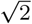 to obtain single-interval sensory uncertainties. This corresponds to the assumption that the subject judges the difference between the stimuli observed in the two intervals. Although this assumption is debated and empirical deviations in both directions were observed (Yeshurun et al., 2008), as discussed in the main text, this assumption is a natural starting point when converting between 2IFC and single-interval perceptual uncertainties. Because the stimuli in the 2IFC task were shown for 15 frames and the computational models of the tracking task operate on single frames, we apply a second correction factor to the perceptual uncertainty estimates. Assuming optimal integration of the Gaussian uncertainty across frames, the 2IFC sensory uncertainty estimates were multiplied by a constant factor of 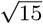.

### G.2 Motion tracking

We reanalyze the data from experiment 2 of Bonnen et al. (2017), in which subjects tracked a circular target on a gray background with their cursor. Instead of the computer mouse, the cursor was controlled used a Leap Motion hand tracking device (Leap Motion, San Francisco, USA). There were four different standard deviations for the target’s random walk. Each subject completed 20 trials per standard deviation in randomly interleaved blocks of 10 trials. As above, the response time series were shifted by 12 time steps w.r.t. the target time series.

### G.3 Gaze tracking

We reanalyze the data from an experiment on gaze tracking (Knöll et al., 2018), in which two macaques (M1 and M2), one human participant (H), and a marmoset (C) tracked the center of an optical flow field displayed on a screen, while their eye movements were recorded. In this experiment, the temporal shift applied to the response time series was 10 time steps, which was determined from the CCGs.

## H Supplementary results

### H.1 Cross-validation

In addition to the model fits on the data from all conditions, we performed a cross-validation evaluation for the tracking experiment with different random walk conditions Bonnen et al., 2017, Appendix G.2. Specifically, we fit all models to the data in a leave-one-condition-out cross-validation scheme, i.e. we fit the model on data from four conditions and evaluate it on the remaining condition, and repeated this procedure for each condition. As a metric for evaluation, we compute the log predictive posterior density relative to the best performing model (Δlog*p*). It was higher for the subjective model than for the other models in all but one condition of one participant (Figure S4 A). To check which model yields the most reliable estimates of perceptual uncertainty, we look at the posterior mean estimates of *σ* (Figure S4 B). The subjective model’s estimates are most consistent across cross-validation conditions, with an average standard deviation of 0.25 arcmin, compared to 0.60 for the basic LQG and 0.52 for the KF. This suggests that the model recovers the subjects’ perceptual uncertainty more reliably than the alternative models.

### H.2 Application to eye tracking data

We reanalyze data from the experiment described in Appendix G.3. Since the generative model of the target’s random walk was slightly different compared to the other experiments, the model was adapted accordingly.

The target dynamics has a velocity component which depends on the current position, such that the target tends towards the center of the screen. The matrices of the dynamical system and cost function are

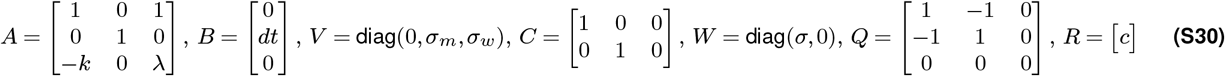

Thus, the agent’s estimate is also three-dimensional, i.e. the agent estimates a velocity from their observations of the position.

As for the other experiments, we fit all four models to the data and computed WAIC as a model comparison metric (Figure S6). Again, the behavioral data is better accounted for by the LQG models compared to the KF model. As in the other analyses of experimental data, the bounded actor models with internal costs better account for the data than the optimal actor model.

## Supplementary figures

**Figure S1.**
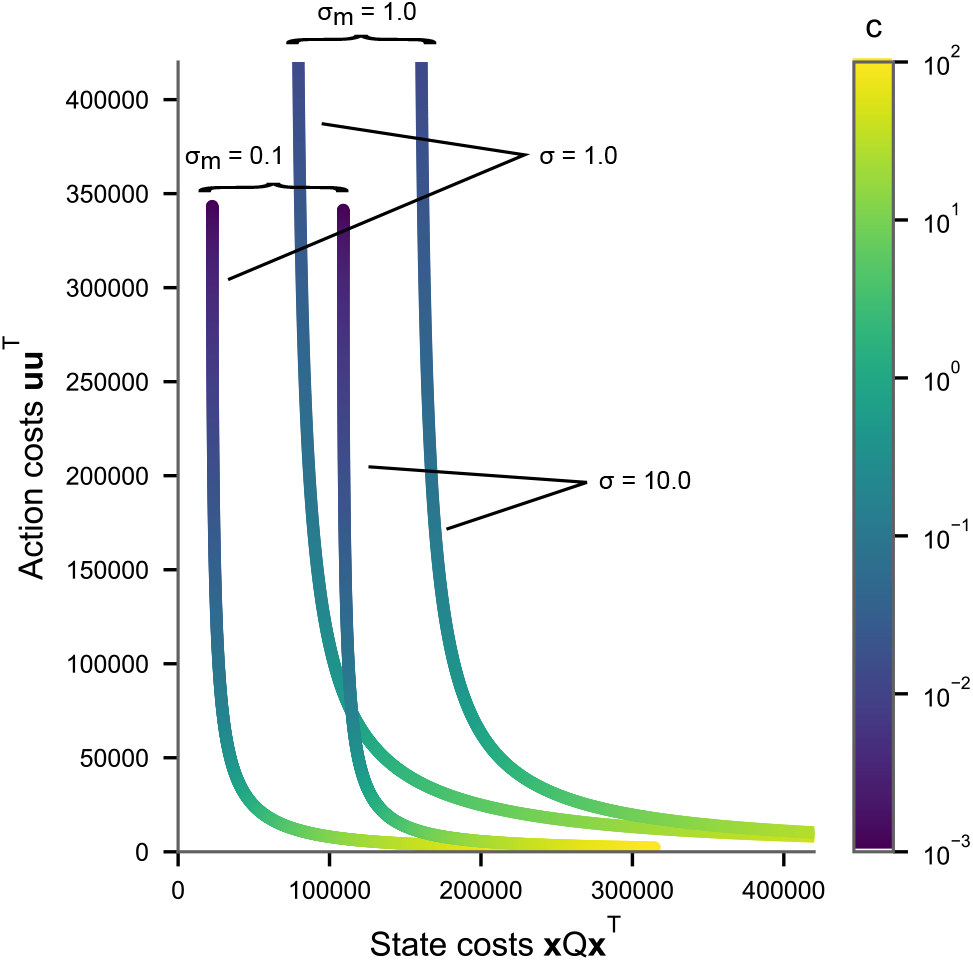
Pareto efficiency plot. Relationship between the two terms of the cost function, i.e. state costs **x***Q***x**^*T*^ and action costs **uu**^*T*^, for different values of the control cost parameter *c*. For the bounded actor model, there is a trade-off between tracking the target (state costs) and expending control effort (action costs). For a specific level of action variability *σ_m_* and perceptual uncertainty *σ*, there is a minimum level of state costs incurred, which can not be alleviated by decreasing action costs via *c*.

**Figure S2.**
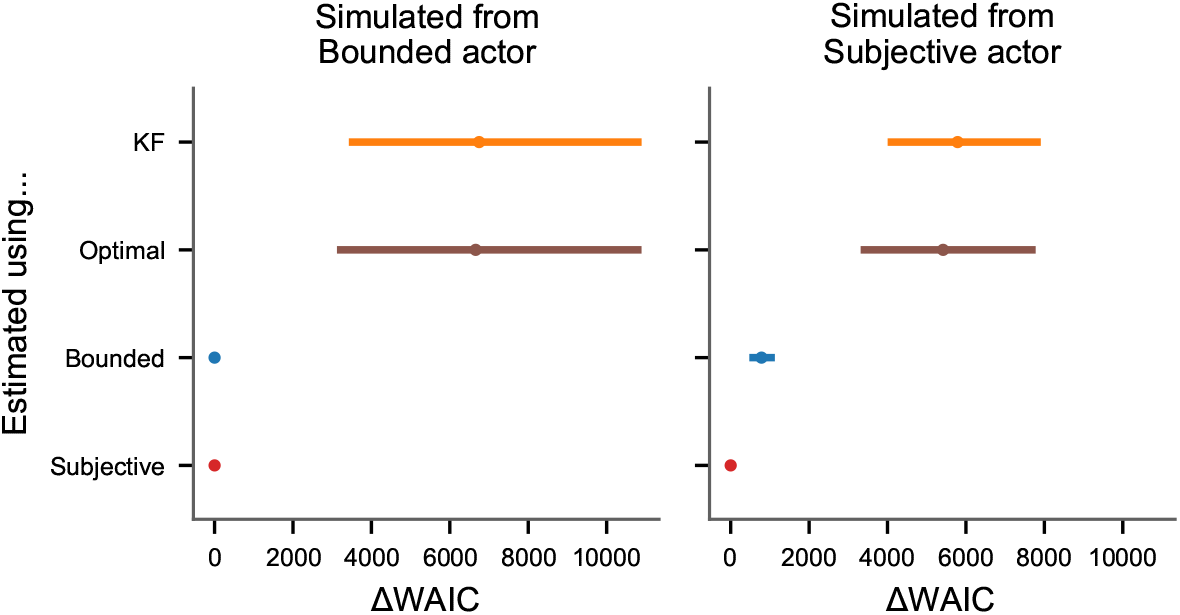
Model comparison for simulations. Model comparison using WAIC between different models fit to data simulated from the bounded actor or subjective actor model.

**Figure S3.**
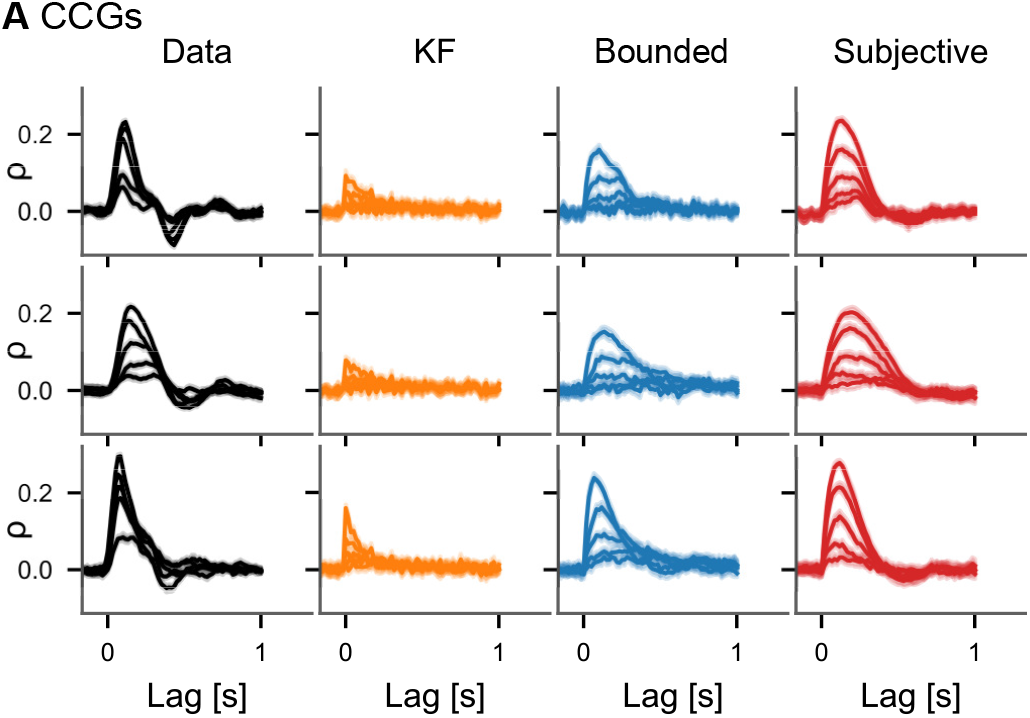
Cross-correlograms. CCGs as in Figure 5 for all three subjects separately based on the data in Bonnen et al. (2017).

**Figure S4.**
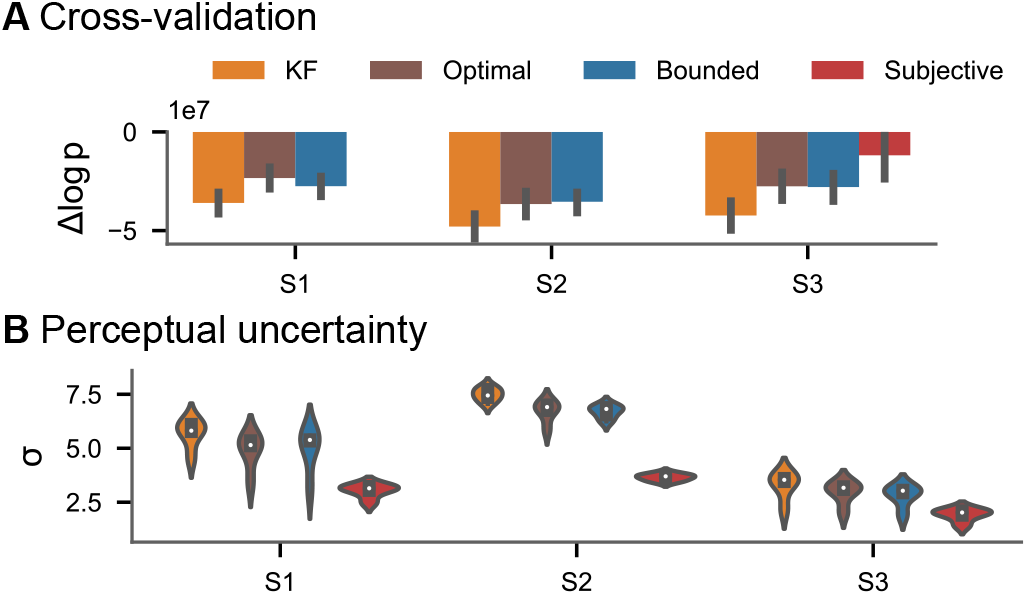
Cross-validation of estimated parameters. **A** Difference in log-posterior w.r.t. worst best per left-out condition (each model was fitted on the remaining four conditions) from Bonnen et al. (2017). In all but one condition of one participant, the model with a subjective component beats the other models. **C** Perceptual uncertainty (posterior mean) estimates across cross-validation conditions.

**Figure S5.**
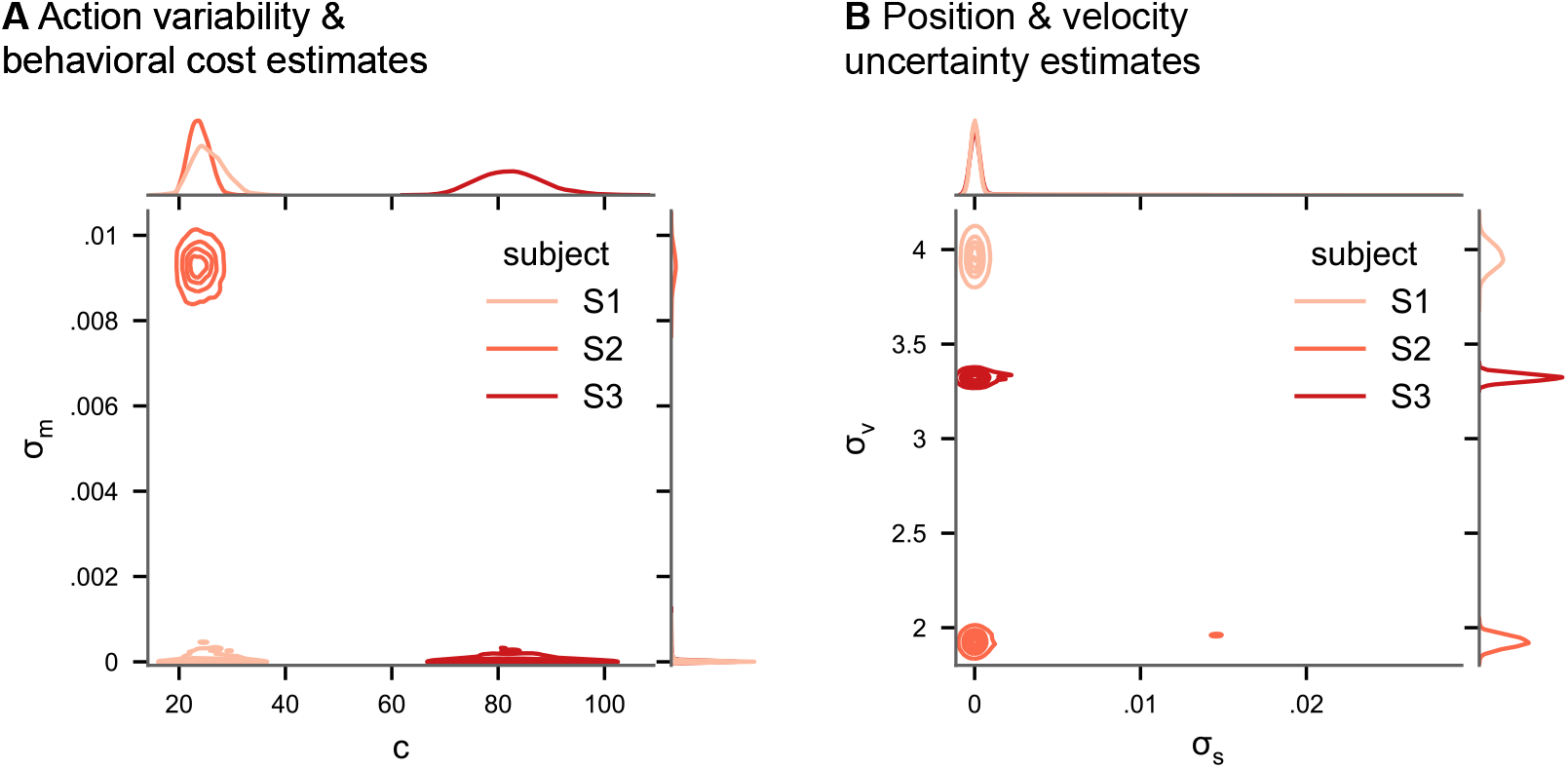
Posterior distributions of model parameters of the subjective model in experiment 2. **B** Posterior distributions for action cost (*c*) and action variability (*σ_m_*). **C** Posterior distributions for subjective stimulus dynamics parameters (position: *σ_s_*, velocity: *σ_ν_*).

**Figure S6.**
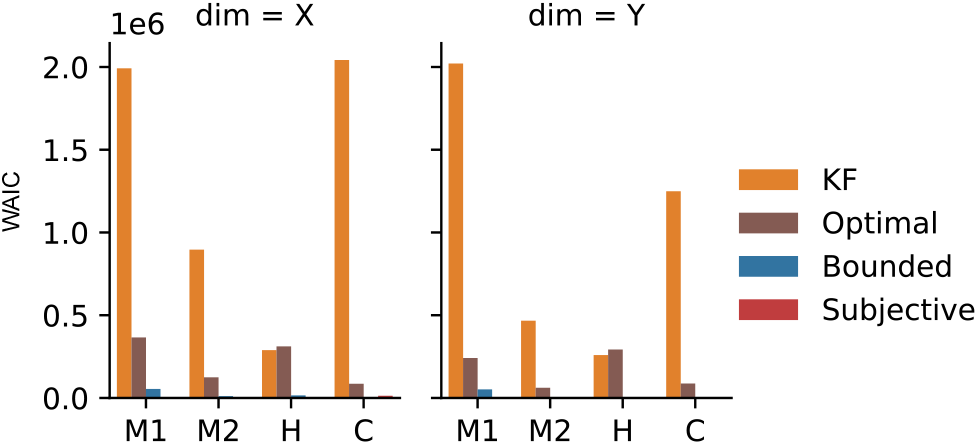
Model comparison on eye-tracking data. WAIC on our four models for the eye-tracking data, fit separately to the X and Y dimension.

